# A public resource of 15 genomically characterised representative strains of *Shigella sonnei*

**DOI:** 10.1101/2025.02.03.635664

**Authors:** Sydney L. Miles, Jane Hawkey, Ben Vezina, Vincenzo Torraca, Claire Jenkins, François-Xavier Weill, Stephen Baker, Kate S. Baker, Serge Mostowy, Kathryn E. Holt

## Abstract

*Shigella sonnei* is rapidly emerging as the dominant agent of shigellosis, an enteric disease responsible for a significant burden of morbidity and mortality worldwide. Whole-genome sequencing of *S. sonnei* isolated over the last three decades has revealed phylogenomic diversity within the population and the emergence of multiple lineages associated with distinct epidemiological patterns such as resistance to critical antimicrobials and/or transmission within different groups. However most experimental work on *S. sonnei* biology and pathogenicity has focused on a single laboratory strain (53G), which is phylogenetically distant from currently circulating strains. Here we introduce a set of 15 phylogenetically diverse and epidemiologically relevant *S. sonnei* isolates made available through publicly accessible culture collections as a resource for laboratory science. We present their complete whole-genome sequences, including the pINV invasion plasmid (missing from a large proportion of public genome data due to loss during laboratory culture). Finally, the characterisation and comparison of these complete genome sequences highlight evidence for ongoing adaptive evolution in *S. sonnei*, featuring the accumulation of insertion sequences, gene pseudogenisation and structural variation.

**Significance as a BioResource to the community:** Genomic analysis of *Shigella* has historically been challenging due to presence of hundreds of repetitive sequence elements (which can cause fragmented assemblies) and loss of the pINV invasion plasmid (essential to virulence) during laboratory culture. Furthermore, most experimental work on *S. sonnei* pathogenicity uses a lab strain that is phylogenetically distant from circulating isolates. To support *S. sonnei* experimental and *in silico* research and increase its relevance to current clinical problems, we report here the complete, high-quality genome sequences of 15 *S. sonnei* isolates, each selected to represent distinct sub-clades of epidemiological interest. We also make the corresponding strains publicly available in national reference culture collections.

**Data summary:** All sequencing reads and complete assemblies have been deposited into the National Center for Biotechnology Information (NCBI) database (accessions to be determined). Genome assemblies and Bakta annotations used in the analysis can be found in Figshare (https://doi.org/10.6084/m9.figshare.28302986) together with Mauve multiple-sequence alignments for the chromosome and pINV plasmid sequences, and genome-scale metabolic models produced for each strain.

Pure cultures of all strains were deposited in the publicly accessible National Collection of Type Cultures (NCTC, UK) or the “*Collection de l’Institut Pasteur*” (CIP, France) (accessions to be determined).

## Introduction

*Shigella* are human-adapted lineages of *Escherichia coli* which have evolved to cause severe diarrhoeal disease, known as shigellosis. Shigellosis is the second leading cause of diarrhoeal deaths globally, most of which occur in children under five years of age in low- and middle-income countries (LMIC) [1].

Within *Shigella*, there are four recognised subgroups that are separated based on their antigenic properties: *S. boydii*, *S. dysenteriae, S. flexneri* and *S. sonnei*. Each *Shigella* subgroup emerged from *E. coli* following the acquisition of a large virulence plasmid (pINV, 210-240 Kbp) which conferred the ability to invade human cells [2, 3]. Aside from pINV acquisition, several other gain- and loss-of-function events have been established as stepwise evolutionary changes which facilitated the human adaptation of *Shigella* [4, 5]. Throughout the course of pathoadaptation, considerable gene loss has been observed [5], a process that is also documented in other host-restricted pathogens such as *Salmonella enterica* serovar Typhi*, Bordetella pertussis* and *Mycobacterium leprae* [6–8]. In the case of *Shigella*, gene loss is mostly associated with insertion sequences (ISs), small transposable elements that can mobilise within the genome, disrupt coding sequences and mediate genome rearrangements, insertions and deletions [5, 9]. Interrogation of *Shigella* and *E. coli* genomes has revealed that each *Shigella* subgroup harbours significantly more copies of ISs than other *E. coli* pathotypes [5, 10].

*S. sonnei* represents the youngest and least genetically diverse *Shigella* subgroup [11], with multidrug-resistant clones emerging as the dominant agent of shigellosis in both high-income and economically transitioning countries [12–14]. The current population of *S. sonnei* is delineated into five lineages, each with varying degrees of global dissemination and expansion [15]. Lineage 1 represents an ancestral lineage rarely detected outside of Europe; Lineage 4 represents an extinct lineage comprising of a single known isolate [11]; Lineage 5 is restricted to Latin America and parts of Africa [16]. Lineage 2 has undergone limited dissemination, establishing localised clones in some regions, but overall, Lineage 3 has been the most successful at global dissemination having been detected on every continent, and today represents the most epidemiologically significant lineage [15]. Within Lineage 3, clades 3.6 (Central Asia III/CipR) and 3.7 (Global III) dominate the epidemiological landscape, with regional studies highlighting a pattern of clonal replacement and expansion, seemingly driven by the independent acquisitions of genes conferring multidrug resistance (encoded on a class II integron (Int*2*)-bearing transposon, Tn*7*) [13, 17].

Increased resistance to antimicrobials is a clear signature of Lineage 3 *S. sonnei*, but there is evidence of ongoing adaptation elsewhere in the genome, marked by the expansion of ISs and the loss of catabolic genes [10, 11]. However, a lack of completed genomes (and even fewer genomes containing pINV [11], which is readily lost during laboratory culture due to the deletion of toxin-antitoxin systems involved in plasmid maintenance [18]) has left gaps in our knowledge of IS-mediated diversity and genome variability. Furthermore, genomic data reveals that the classical laboratory strain 53G (isolated in 1954 and used for the vast majority of experimental work on *S. sonnei* [19]) belongs to Lineage 2 which is now comparatively rare among clinical isolates and is quite distant from the dominant Lineage 3 [11, 15].

Here we present the complete genomes of 15 pINV-containing *S. sonnei* isolates, including representatives from lineages 1, 2 and 3. The comparison of completed genomes supports ongoing adaptive evolution in Lineage 3, characterised by increased IS abundance, gene pseudogenisation and genomic rearrangements. Overall, the results presented here provide novel insights into the genomic variation of *S. sonnei* lineages and significantly further the resources available for the study of *S. sonnei*.

## Methods

### Bacterial strains used in this study

*S. sonnei* isolates used in this study are listed in **Tables 1 and S1**. Isolates were originally collected as part of routine public health surveillance in the United Kingdom [20] and France [21], or as part of a cohort study in Vietnam [22]. Strains were received on agar slants or stabs and streaked Tryptic Soy Agar (TSA) plates, supplemented with 0.01% Congo Red to select for pINV+ isolates [23]. Smooth, red colonies (which indicate T3SS and pINV presence) were picked and stored in 25% (v/v) glycerol at - 80°C.

### Genome sequencing

The DNA extraction, long-read (Oxford Nanopore Technology) and short-read (Illumina) sequencing of *S. sonnei* clinical isolates were performed by MicrobesNG as a paid service. Briefly, an overnight culture was grown in tryptic soy broth ((TSB), incubation at 37°C, 400 rpm) and sub-cultured to prepare isolates for sequencing. Once the sub-culture reached mid-exponential phase, cells were pelleted by centrifugation at 4000 *x g* for 4 minutes, washed in 1 mL PBS and resuspended at a density of 4x10^9^ cells in 500 μL DNA shield (Zymo Research). For Illumina sequencing, paired-end libraries were prepared with the Nextera XT Library Prep Kit and sequenced on an Illumina NovaSeq 6000 instrument to generate 250 bp reads. For ONT sequencing, libraries were prepared with the SQK- RBK114.96 kit (Oxford Nanopore Technologies) and loaded onto an R.10.4.1 flow cell for sequencing on a GridION instrument and basecalled using Dorado (model r1041_e82_400bps_hac_v4.2.0).

### *De novo* genome assembly

Prior to genomic analysis, sequencing reads were quality checked using FastQC (v.0.12.0) [24], trimmed using Filtlong (v.0.2.1) [25] (for long reads) or Trimmomatic (v.0.4.0) [26] (for short reads). Genomes were then assembled using the Hybracter (v.0.7.3) [27] long-read first assembly pipeline, with Flye (v.2.9.4) [28] selected as the long read assembler. As part of the pipeline, genomes were polished with Medaka (v.1.8.0) first, then with short reads using PyPOLCA (v.0.3.1) [29, 30]. The quality of assemblies was checked using Quast (v.5.0.2) [31] (using 53G as a reference genome (accession NC_016822)). Completeness and contamination were analysed using CheckM (v.1.2.3) [32]. Contigs were concatenated into a multifasta file using SeqKit (v.2.8.2) [33] and then genomes were annotated using Bakta (v.1.9.3) [34], with the full database option selected.

### Genome characterisation

Complete genomes were genotyped using the *S. sonnei* genotyping framework implemented in Mykrobe (v.0.13.0) [15, 35], to confirm *S. sonnei* lineage assignments matched those obtained previously from published Illumina sequence data for the same isolates [15]. Plasmid sequences were replicon-typed using MOB-suite (v.3.1.9) [36] ‘Mob-typer’ function. Antimicrobial resistance (AMR) determinants were identified using NCBI AMRFinderplus (v.3.12.8) [37]. Colicins were identified using ABRicate (v.1.0.1) [38] using the custom colicin database created by De Silva *et al* [39] (https://figshare.com/articles/dataset/colicin_database/20768260/1?file=37009930).

IS elements were identified using ISEScan (v.1.7.2.3) [40] and results were filtered to identify ISs present in the chromosome, pINV and other plasmids. Bakta did not efficiently annotate pseudogenes (tested on 53G for which the number of pseudogenes was previously reported [5, 10]), so the prokaryotic genome annotation pipeline (PGAP) (v.6.7) [41] was used to quantify and compare pseudogenes.

### Pangenome analysis

Pangenome analysis was performed using Panaroo (v.1.5.0) [42] using the ‘strict’ mode, MAFFT (v.7.526) [43] was selected to perform core genome alignment, and otherwise default settings were used. The phylogenetic tree for pangenome visualisation was produced using the core genome alignment file generated by Panaroo as an input to FastTree (v.2.1.1) [44] which was run using the generalised time-reversible model and otherwise default settings. Visualisation was performed using Phandango (accessed on 18/08/2024) [45]. Unique HGCs for each lineage were classified into biological process Gene Ontology (GO) categories using the PANTHER functional classification system (v.19.0) [46].

### Whole genome alignment

Complete genome sequences were reoriented to the *fabB* gene where necessary due to an inverted colinear block involving the *dnaA* gene. Whole genome alignments were performed using progressive Mauve (v.2.4.0) [47]. This approach generates locally colinear blocks which enables the visualisation of structural genome rearrangements. To create isolated gene cluster comparison figures, relevant gene clusters were extracted following visualisation in Mauve and aligned using Clinker (v.0.0.29) [48].

### Genome-scale metabolic models

Metabolic models were constructed using CarveMe (v1.5.1) with the following options: ‘-u gramneg -g M9’ [49]. Metabolic reaction presence/absence was generated using reaction_presence_absense_generate.py (https://github.com/bananabenana/Metabolic_modelling_scripts/tree/main/reaction_pres_abs). Growth simulations were then performed with Bactabolize (v1.0.3) using the ‘fba’ command [50].

## Results and discussion

### The complete genomes of 15 phylogenetically diverse *S. sonnei* isolates

*S. sonnei* isolates were selected based on existing short-read (Illumina) data to represent the diversity of the current *S. sonnei* population. They comprised two Lineage 1 isolates, four Lineage 2 isolates (including 53G) and 10 Lineage 3 isolates (**Table S1**). We initially sought to include representatives from Lineage 4 and 5 but found that cultures were either non-viable following many years of laboratory storage or simply had lost pINV (including the only known Lineage 4 isolate) and thus were unsuitable for inclusion. Complete chromosomal and pINV sequences were obtained using hybrid short- and long-read sequencing; general features of the newly sequenced isolates are summarised in **Table 1**.

**Table 1.** Summary of *S. sonnei* genome assemblies presented here. ^1^Genotypes were predicted using the *S. sonnei* genotyping framework implemented in Mykrobe, ^2^CDSs and tRNAs were predicted using Bakta. *Features of the completed genome of the classical laboratory strain, 53G, are included for comparative purposes. ONT, Oxford Nanopore Technologies.

| Strain ID | Genotype<sup>1</sup> | Genome size (bp) | ONT depth | CDSs<sup>2</sup> | tRNAs | Contigs | Chromosome size (bp) | pINV size (bp) | Other plasmids |
| --- | --- | --- | --- | --- | --- | --- | --- | --- | --- |
| 201809330 | 1 | 5389660 | 61x | 5428 | 97 | 7 | 4916714 | 242731 | 5 |
| 391324 | 1.5 | 5260824 | 59x | 5255 | 97 | 4 | 4888493 | 238313 | 2 |
| 356538 | 2.1 | 5170588 | 76x | 5176 | 97 | 4 | 4946399 | 216858 | 2 |
| 830292 | 2.3 | 5150027 | 60x | 5166 | 96 | 3 | 4924717 | 220157 | 1 |
| 373220 | 2.12.4 | 5289501 | 47x | 5306 | 93 | 4 | 4983563 | 223662 | 2 |
| 590907 | 3.4.1 | 5174681 | 74x | 5229 | 97 | 7 | 4940164 | 215149 | 5 |
| 623218 | 3.6.1 | 5112917 | 68x | 5154 | 97 | 9 | 4862817 | 214316 | 7 |
| 02_1157 | 3.6.1.1.1 | 5073758 | 86x | 5110 | 96 | 5 | 4832017 | 212976 | 3 |
| 642321 | 3.6.2 | 5100173 | 43x | 5135 | 97 | 6 | 4866958 | 213229 | 4 |
| 633497 | 3.7.11 | 4986415 | 46x | 5046 | 93 | 7 | 4657310 | 214646 | 5 |
| 598955 | 3.7.16 | 5039937 | 54x | 5067 | 98 | 7 | 4807305 | 214536 | 5 |
| 618335 | 3.7.28 | 5138354 | 61x | 5177 | 98 | 6 | 4806102 | 214666 | 4 |
| 03_0142 | 3.7.29.1.4 | 5156461 | 86x | 5182 | 96 | 9 | 4824935 | 214579 | 7 |
| 627346 | 3.7.30.1 | 5122274 | 42x | 5168 | 100 | 6 | 4884953 | 216152 | 4 |
| 381259 | 3.7.30.4.1 | 5260132 | 58x | 5322 | 91 | 8 | 4933886 | 214700 | 6 |
| 53G* | 2.8 | 5220473 | NA | 5248 | 96 | 5 | 4988504 | 215774 | 3 |

There was some variation in genome size, with chromosome sizes ranging from 4.65 to 4.98 Mbp. In line with expectations from prior reports on genome reduction [10], Lineage 1 genomes had an average chromosome size of 4.90 Mbp (range 4.89 to 4.92 Mbp), Lineage 2 chromosomes averaged at 4.96 Mbp (range 4.92 to 4.99 Mbp), whilst Lineage 3 chromosomes were smallest, averaging at 4.84 Mbp (range 4.66 to 4.94 Mbp). Likewise, pINV sizes ranged from 212 to 242 Kbp, and pINV in Lineage 1 genomes was ∼ 24 Kbp larger than Lineage 2 and 3 genomes (average 240 Kbp vs 216 Kbp).

In addition to the chromosome and pINV, each genome also harboured additional plasmids (between 1 and 8 per genome, ranging from 1,459 to 108,503 bp) (**Fig 1A, Table S2**). Plasmid typing using MOB-suite revealed a diverse plasmid repertoire, with Col plasmids (colicinogenic plasmids which encode the genes required for colicin production [51]) being frequently detected. The carriage of colicins E3 and E1 were most common (**Fig 1B, Table S3**) and have associated with local lineage replacement in *S. sonnei* [13, 39], although no global lineage-associated trends were evident amongst the selected isolates.

**Figure 1.**
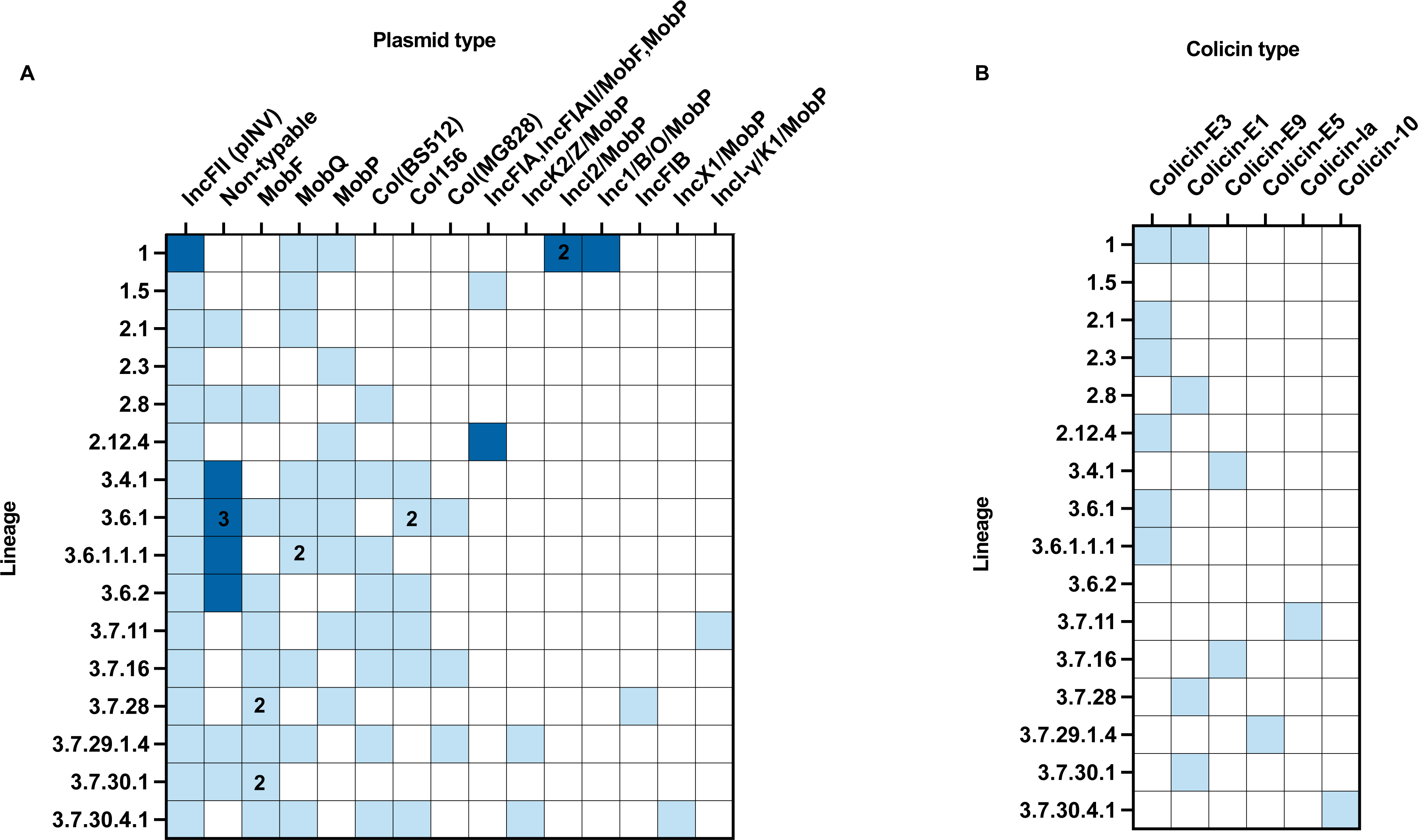
Summary of plasmid content and colicin presence in *S. sonnei* assemblies presented here. **A)** Plasmids were typed using the MobTyper function in MOB-suite. White squares indicate the absence of the plasmid type, light blue indicates its presence, and dark blue indicates that the plasmid type is present and encodes antimicrobial resistance determinants. Where a number is printed in the cell, this indicates multiple of the same plasmid type present in a single strain. **B)** Colicins were identified using ABRicate and the full colicin database created previously [39]. White indicates the absence of colicin type, and light blue indicates its presence.

### Carriage of AMR determinants is representative of known lineage associations

Screening assemblies for known AMR determinants confirmed that the isolates we selected were representative of the AMR profiles that had previously been associated with the genotypes they were selected to represent. Isolates belonging to clade 3.6.1 were both found to carry the GyrA-S83L mutation associated with this sub-clade [52]; the isolate representing subclade 3.6.1.1.1 also carried the additional mutations GyrA-D87G and ParC- S80I, which are characteristic of subclade 3.6.1.1 and its descendants, and are responsible for its high-level fluoroquinolone resistance [53]. All clade 3.6 isolates carried Tn*7* harbouring *dfrA1* and *sat2* in the integron cassette (conferring resistance to trimethoprim and streptothricin), which was inserted between an IS*4*-family transposase and *glmS*. Similarly, all clade 3.7 isolates carried a distinct Tn*7* variant inserted at the same chromosomal locus **(Fig S1A)** harbouring *dfrA1, sat2* and *aadA1* (conferring aminoglycoside resistance). Additionally, all isolates belonging to clade 3.6 were found to carry the small spA plasmid, encoding *tetA*, *aph(6)-Id, aph(3”)-Ib* and *sul2* (**Table 2, Table S3**), as is commonplace within this clade [52]. Genotypes 2.1 and 3.4.1 were both found to harbour distinct variants of the chromosomally located *Shigella* resistance locus (SRL) which encodes for *aadA1*, *tetB*, *catA1* and *bla*_OXA-1_, typical of Latin American-associated *S. sonnei* (**Fig S1B**) [54]. Insertions of the SRL occurred at different sites in the chromosome, but in both cases, they were inserted into a copy of *trnS* (which encodes for tRNA-Ser), consistent with previous reports [55]. Until now, the location of some resistance determinants as either chromosomally encoded (and so stably fixed) or plasmid encoded (which may be more easily lost), was unclear due to a lack of completed genomes. Resolution of the location of resistance determinants may be beneficial for future use in experimental work, where stable chromosomally encoded genes may act as selective markers.

**Table 2.**
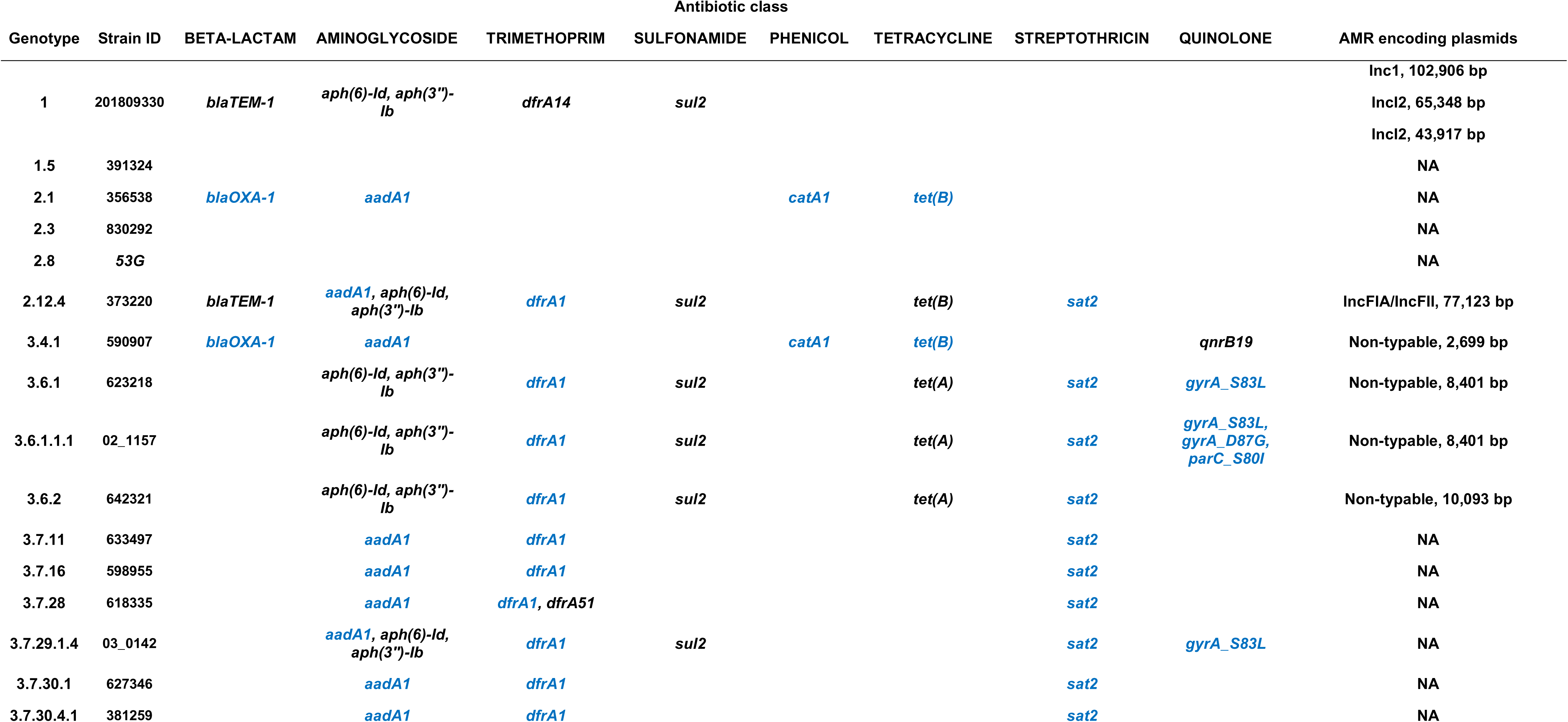
Summary of antimicrobial resistance determinants identified by AMRfinder. Plasmid-encoded determinants are displayed in black, and chromosomally encoded determinants are shown in blue. The column titled ‘AMR encoding plasmids’ indicates the type (as indicated by MOB-suite), sizes and accessions of plasmids encoding antimicrobial resistance determinants in each strain. NA=not applicable, no AMR encoding plasmids identified in this strain.

### Pangenome analysis highlights limited lineage-associated gene content variation

To explore overall gene content variation within *S. sonnei* lineages, a pangenome analysis was performed using Panaroo, including the 15 novel isolates and the laboratory strain 53G. The resulting pangenome was found to be open (γ=0.07) and a total of 6317 homologous gene clusters (HGCs) were identified: of these, 4282 genes were present in all 16 genomes, representing the core genome; 1268 genes were present in ≥3 strains, representing ‘shell’ genes and 767 genes were present in <3 strains representing the ‘cloud’ genes (**Fig 2**). The number of unique HGCs varied from 1-104 per isolate (**Table 3**). Lineage 1 had the most lineage-specific HGCs (HGCs present in every isolate of that lineage and absent in the other lineages) at 63, consistent with previous reports of gene loss in lineages 2 and 3 compared to Lineage 1 [10]. Lineage 2 had only 6 lineage-specific HGCs and Lineage 3 had 27 lineage-specific HGCs. Many lineage-specific HGCs in lineages 2 and 3 were annotated as ISs or IS accessory genes (discussed below). These results highlight minimal fixed gene content variation between *S. sonnei* lineages, suggesting that variation in the success of lineages is not largely driven by gene content alone.

**Figure 2.**
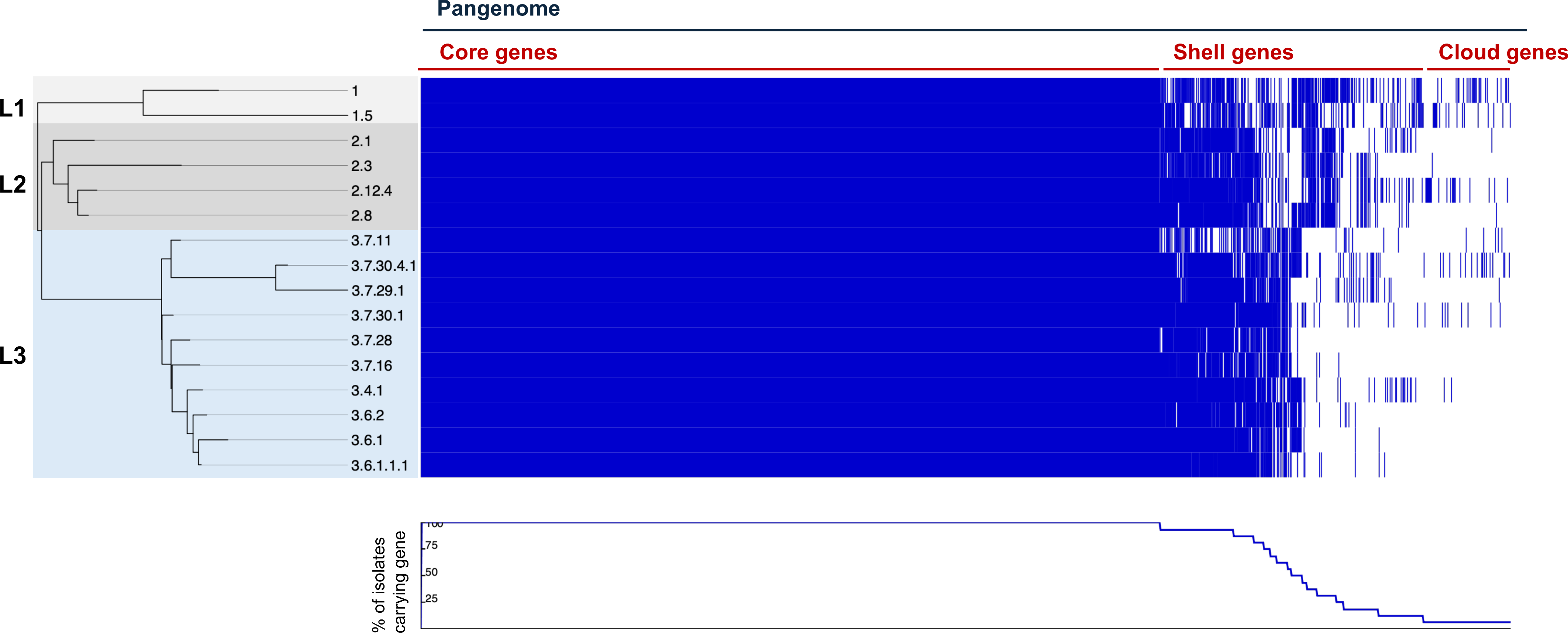
Linear visualisation of the *S. sonnei* pangenome plotted alongside a maximum-likelihood phylogenetic tree. Panaroo was used to build a pangenome and a core genome alignment. FastTree was used to construct a maximum likelihood tree, which is rooted at the midpoint. The resulting outputs were then visualised using Phandango. Blue stripes indicate the presence of a gene, and white stripes indicate the absence of a gene. Core genes are those present in all genomes; shell genes are defined by the presence in ≥3 genomes; cloud genes are those present in <3 genomes.

**Table 3.**
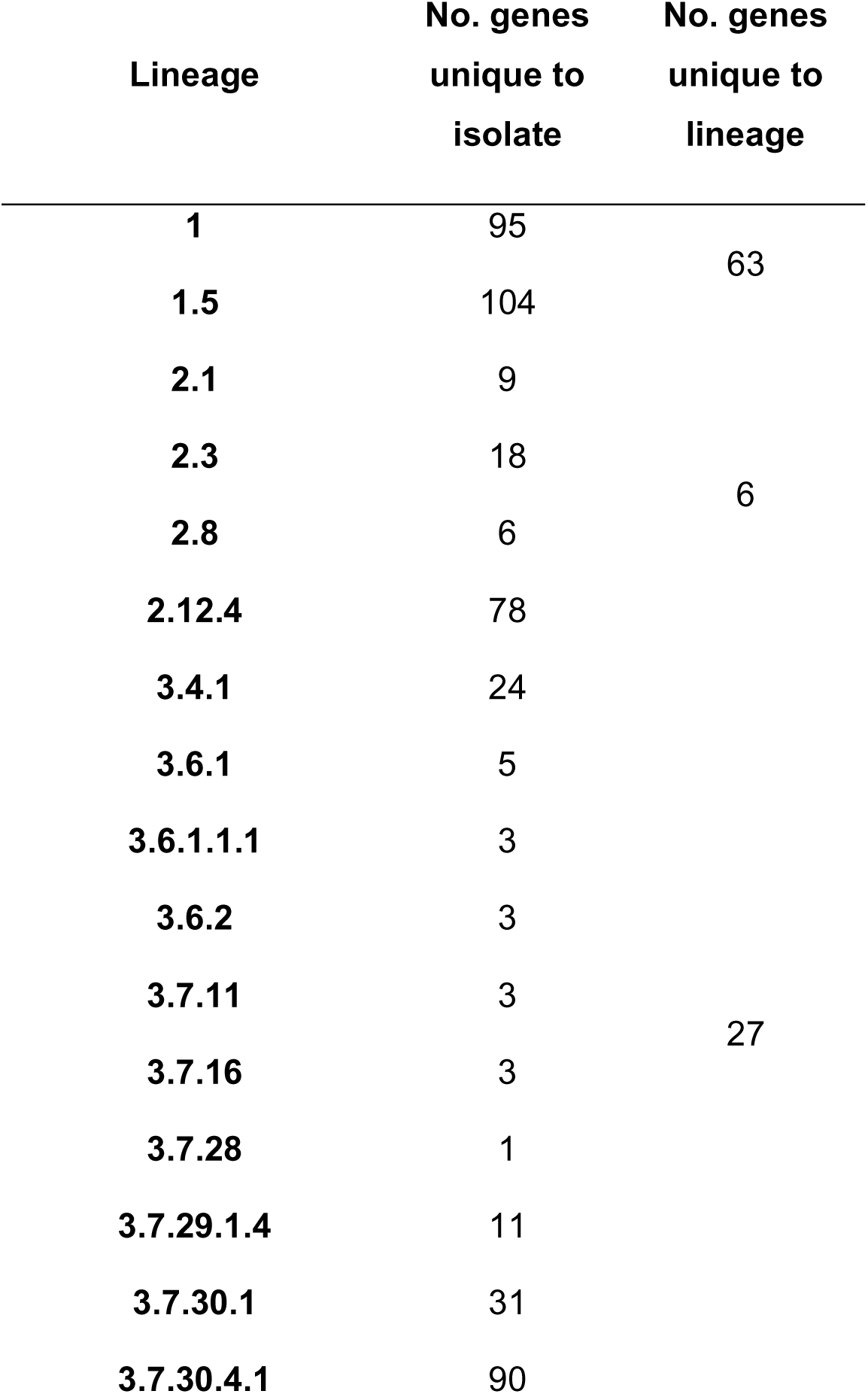
Number of homologous gene clusters (HGCs) unique to each isolate and to each lineage. Unique genes were annotated using Bakta and HGCs were identified using Panaroo.

### Variations in the carriage of insertion sequences

Previous studies have produced evidence for ongoing IS activity within *S. sonnei*, with data from short-read sequences highlighting the proliferation of ISs specifically within lineages 2 and 3 [10]. However, this has not been examined in completed chromosomal sequences, and never in pINV. We therefore examined the abundance and composition of ISs across *S. sonnei* lineages using our reference genomes. A total of 16 distinct IS families were identified across all genomes (**Fig 3A**). There was some variation in the total IS burden by lineage, with Lineage 1 genomes harbouring the fewest, with an average of 480; Lineage 2 had an average of 490 whilst Lineage 3 genomes had an average of 501.

**Figure 3.**
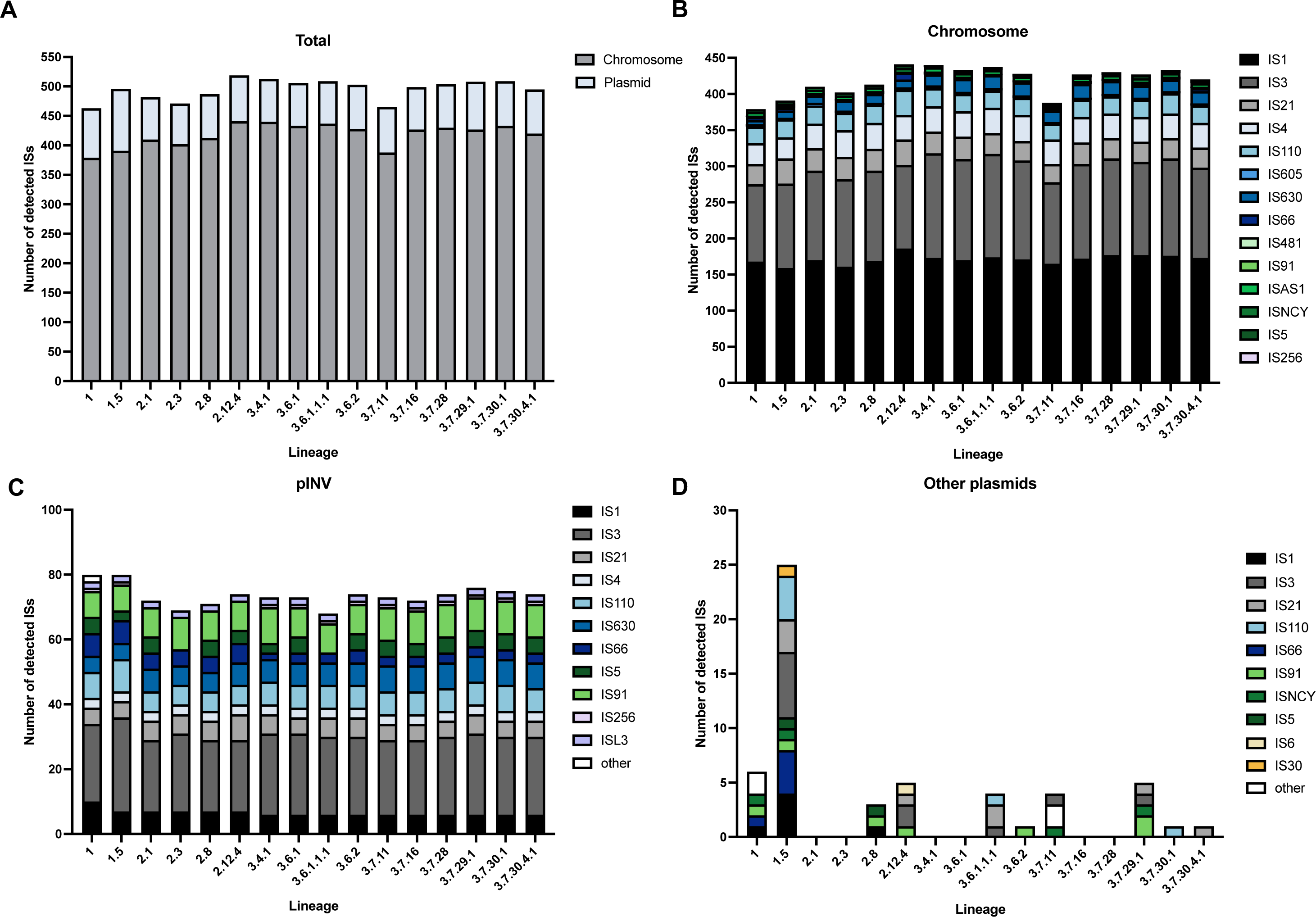
Insertion sequences (ISs) detected in complete *S. sonnei* genome sequences by ISEscanner. **A)** Total number of ISs identified across all contigs of completed genomes and their location. **B)** ISs identified in completed chromosomal sequences. **C)** ISs identified in pINV sequences **D)** ISs identified in other plasmids present in *S. sonnei* genomes. ISs were detected using ISEscanner, ‘other’ represents ISs that could not be matched to a known IS family within the database used.

The detected ISs were split based on their presence in the chromosome, pINV or other plasmids to account for their different evolutionary histories and dynamics. Considering the abundance of ISs in the chromosome alone revealed a clearer trend, with Lineage 1 harbouring the fewest chromosomal ISs (median 385, range 379 to 391) and Lineage 3 harbouring the most (median 429, range 388 to 440) (**Fig 3B**). In pINV, a total of 12 different IS families were identified and two were unique to pINV: IS*5* and IS*L3*, neither of which have been reported in *S. sonnei* before to our knowledge, likely owing to a scarcity of pINV sequences. A contradictory trend in IS abundance in pINV (compared to chromosomal IS abundance) was observed (**Fig 3C**), consistent with pINV being larger in this lineage, although the differences in IS count were smaller compared to chromosomal ISs (median 80 in Lineage 1; median 71.5 in Lineage 2, median 73.5 in Lineage 3). For ISs carried on other plasmids, Lineage 1.5 harboured the most (n=25), which is consistent with its carriage of the largest plasmid (126 Kbp IncFIA,IncFII/ MOB_F_,MOB_P_ plasmid; **Fig 3D**). ISs carried on other plasmids were sporadically detected in single strains, but there was no clear lineage-associated pattern, in line with the limited lineage-associated trends in plasmid carriage.

### Lineage 3 genomes harbour more pseudogenes

The accumulation and proliferation of ISs have been linked to increased formation of pseudogenes (inactivated gene remnants) [56]. To investigate any changes in gene functionality, genomes were screened for the presence of pseudogenes using PGAP. The number of pseudogenes predicted using this method is higher than previously published predictions (for 53G at least [10]), but this is likely due to differences in the annotation method, whereby PGAP may be recording 2 separate pseudogenes in situations where a single gene is interrupted by an IS thus creating 2 fragments.

In agreement with the increased abundance of ISs in Lineage 3 isolates, this analysis also revealed a similar increase in the total number of pseudogenes present in Lineage 3 genomes, with mean 711 identified compared to 671 in Lineage 2 and 663.5 in Lineage 1 (**Fig 4A-D**). The differences in pseudogene number were mostly concentrated in the chromosome, where the Lineage 3 chromosomes harboured the most pseudogenes (**Fig 4B**). In pINV, the same pattern as IS abundance was observed in the number of pseudogenes, where Lineage 1 genomes had the greatest abundance of pseudogenes (**Fig 4C**), likely due to the larger pINV size.

**Figure 4.**
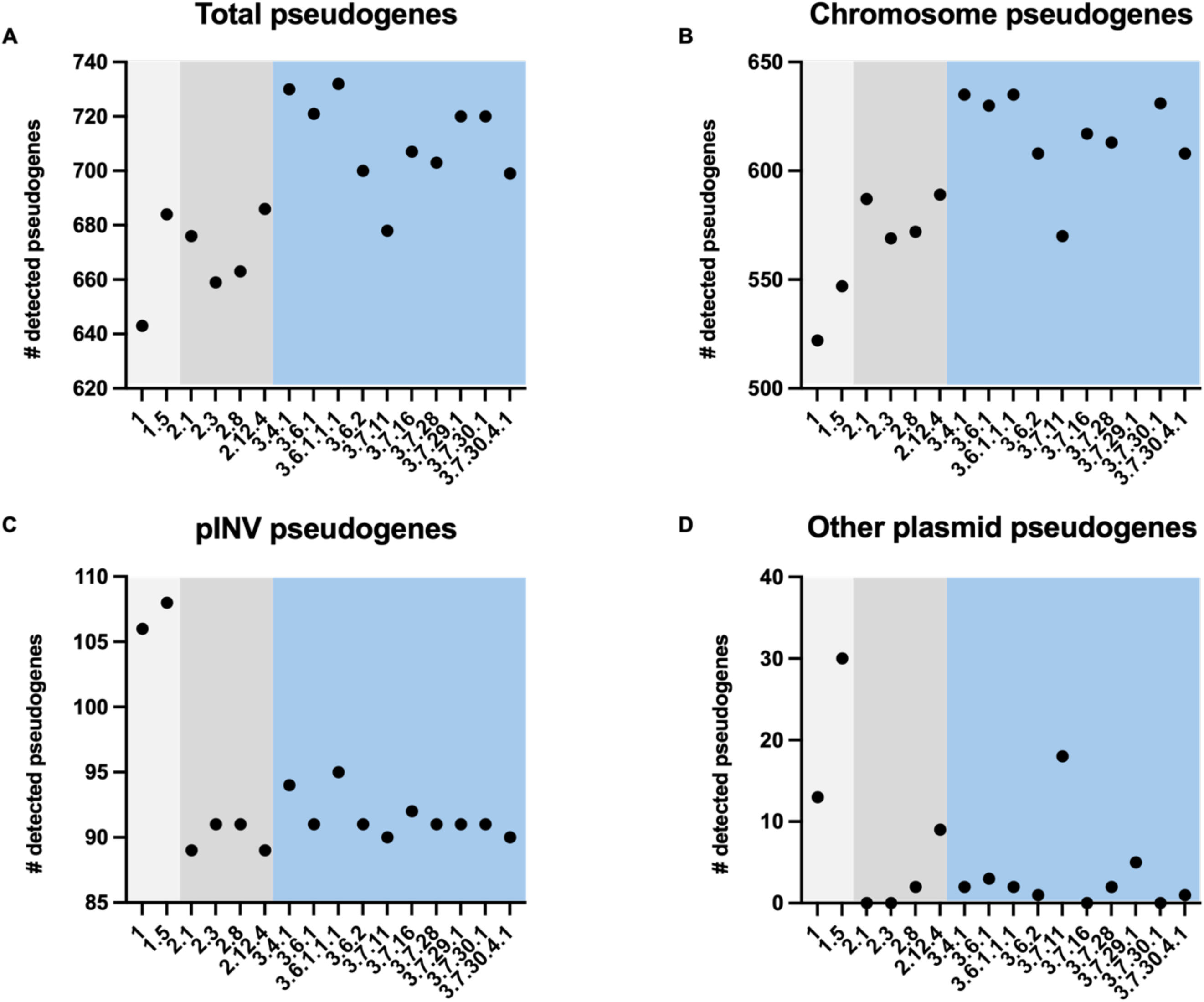
Abundance of pseudogenes predicted by the NCBI prokaryotic genome annotation pipeline (PGAP). **A)** Total pseudogenes identified across all contigs of completed genomes. **B)** Pseudogenes identified in completed chromosomal sequences. **C)** Pseudogenes identified in pINV sequences. **D)** Pseudogenes identified in other plasmids present in *S. sonnei* genomes. Lineage 1 isolates are in light grey, Lineage 2 in dark grey and Lineage 3 in blue.

### Structural variation in *S. sonnei* genomes

*Shigella* genomes are documented to have an exceptionally high rate of structural variation compared to other pathogenic (and non-pathogenic) *E. coli* [57], likely mediated by the large complement of ISs in their genomes [10]. To investigate lineage-associated structural variation, whole-genome alignments of chromosomal and pINV sequences were inspected.

In the chromosomes, we identified four large-scale (>7 Kbp) variations that were conserved within lineages (summarised in **Table 4**) two of which were identified in all Lineage 2 and 3 genomes, and two of which occurred in only Lineage 3 genomes. In lineages 2 and 3, of particular note was the absence of *xylFGH* and *xylR*, which have a role in xylose utilisation [58], which suggests adaptive loss of metabolic function, since these genes are also disrupted in other *Shigella* subgroups [5]. We also observed the complete loss of Type 1 fimbrial operon in all Lineage 3 genomes and distinct partial disruptions of the operon in Lineage 1 and 2 genomes. However, whether this confers a functional difference is unclear. The convergent disruption of type 1 fimbriae has been previously reported in all other *Shigella* subgroups [59], suggesting that its presence is detrimental to *Shigella*, and thus its parallel loss in multiple lineages may reflect adaptive evolution.

**Table 4.**
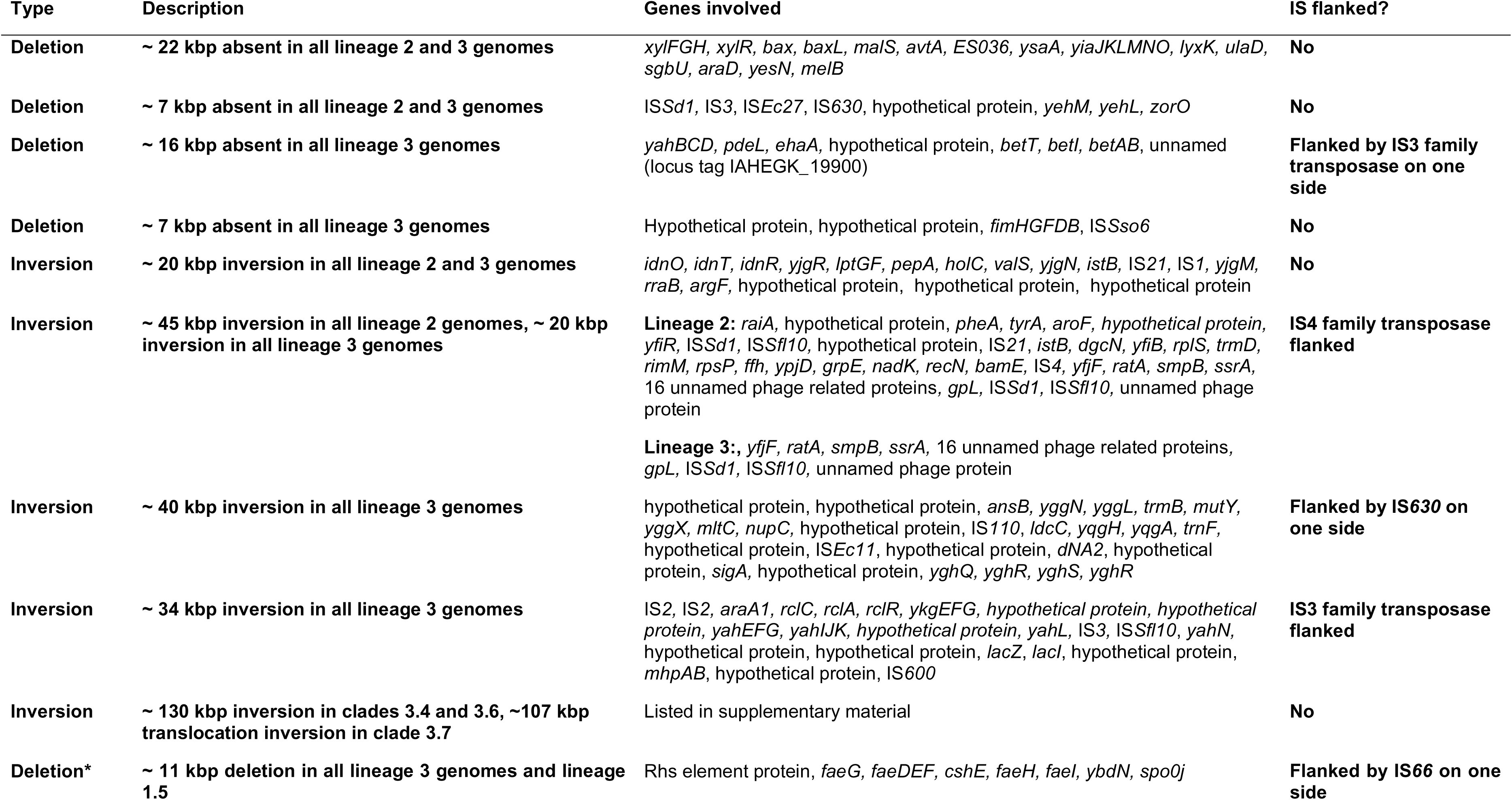
Summary of conserved, large-scale structural variants identified. . Only variants conserved in every isolate of each lineage, and those over 7 Kbp are reported here. *indicates structural variation detected on pINV; all other variations were found to be chromosomal.

We identified five large-scale inversions conserved within lineages. Two were identified in all Lineage 2 and 3 genomes (∼20 and 45 Kbp), and in this case both were flanked by IS*4*- family transposases, suggestive of IS-mediated homologous recombination. Three inverted regions were identified uniquely in Lineage 3 genomes but only one of these was flanked by IS copies, suggesting that other mechanisms may also be driving structural variation in *S. sonnei*. The implications of these chromosomal rearrangements remain ambiguous, but potential impacts on gene expression, or inactivation, are likely important for conferring adaptive changes. Furthermore, lineage-associated variations identified here might indicate differential niche occupation, since rearrangements are known to contribute to bacterial adaptability [60].

For pINV, we observed limited structural variation, and the differences in pINV size (described above) did not seem to be mediated by a single large structural rearrangement, but instead by several smaller-scale deletions. The most prominent example is an ∼11 Kbp loss shared by all Lineage 3 genomes, as well as Lineage 1.5, which included the region encoding for a K88 fimbriae (previously reported to play a role in the pathogenesis of enterotoxigenic *E. coli* [61]). The overall lack of pINV variation observed here is consistent with an essential role for pINV maintenance in the pathogenesis and spread of *S. sonnei* [62].

### Metabolic capacity of *S. sonnei* genotypes

The convergent loss of metabolic function has been previously documented in all *Shigella* subgroups [10, 63], and likely represents a key step towards host-adaptation. To investigate any lineage-associated trends in the metabolic capacity of *S. sonnei*, strain-specific genome-scale metabolic models were built, and growth capacities were simulated. On average, the number of metabolic reactions predicted for each lineage was similar **(Fig 5A, Table S5)**; a total of 2898 metabolic reactions were identified, of these, 2505 were shared between all strains. A median of 2748 reactions were identified in Lineage 1 strains (range 2742 to 2753), 2741 in Lineage 2 (range 2625 to 2746) and 2691 in Lineage 3 strains (range 2602 to 2749). Few reactions were specific to a single lineage: seven were predicted in only Lineage 1 isolates (Alpha amylase, periplasmic alpha-amylase, dextrin import, starch import, Glutathione transport, Succinate dehydrogenase and L-cysteine-specific tryptophanase); two predicted only in Lineage 1 and 2 isolates (Betaine-aldehyde dehydrogenase and a raffinose proton symporter) and four predicted in only Lineage 3 isolates (D-galactose ABC transporter, Glutamate synthase. Loss of a polysaccharide ABC transporter resulted in predicted inability to import maltotetraose, melibiose and raffinose). For growth simulations, no growth was detected under anaerobic conditions on any substrates (despite *Shigella* being a facultative anaerobe), so only aerobic growth was considered here. Consistent with the metabolic reaction data, we identified a median of 410 predicted growth phenotypes in Lineage 1 (range 408-412), 398 (range 384-407) in Lineage 2 and 396 (range 374-422) in Lineage 3. All strains were predicted to grow on 350 substrates, with variability observed for 105 substrates **(Fig 5B)**. The predicted ability to utilise L-Tryptophan was restricted to Lineage 1, which likely represents another instance of adaptive change, since the inability to utilise tryptophan (resulting in the indole-negative phenotype) has been lost in parallel across many *Shigella* lineages [64]. Otherwise, there were no predicted growth phenotypes that were unique to a single lineage, this variation between presence of reactions and predicted growth could be explained by the exclusion of anaerobic growth conditions in our analysis, as well as being unable to model unknown/novel metabolism likely encoded within *Shigella*. This data suggests that there may be an overall trend towards metabolic streamlining in Lineage 3 *S. sonnei*, however, the differences observed here are minor and would benefit from functional verification.

**Figure 5.**
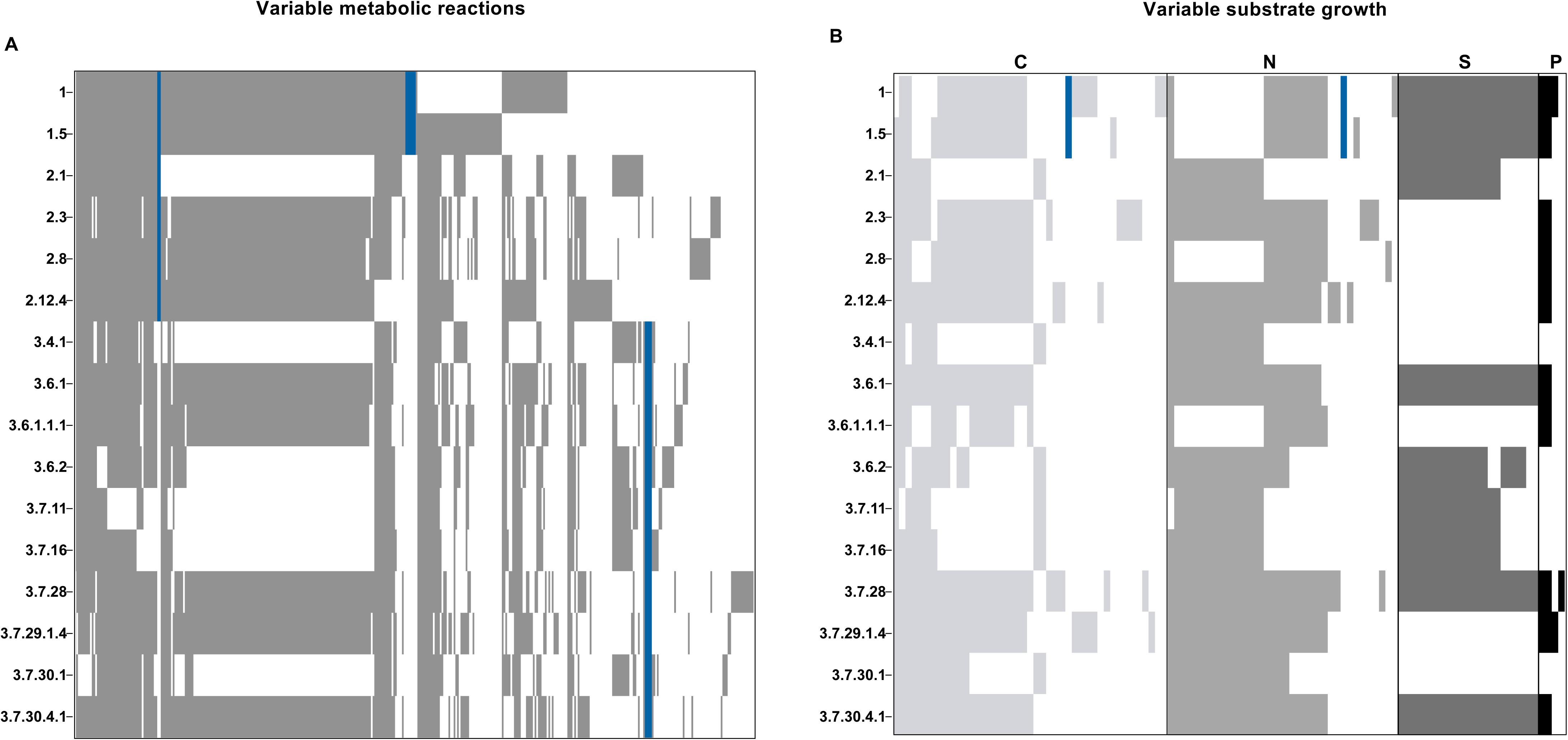
Heatmap of variable metabolic capacities predicted for each *S. sonnei* genome. **A)** Presence or absence of variable metabolic reactions in each strain. Grey indicates presence, white indicates absence and blue indicates reactions unique to one or more lineages, n=394 reactions presented. **B)** Predicted growth simulations on carbon (C), nitrogen (N), sulphur (S) or phosphorous (P) based metabolites. Grey or black indicates presence, white indicates absence and blue indicates reactions unique to one or more lineages, n=105 substrates presented. Full genome-scale metabolic models and tables summarising all metabolic reactions and growth capacities can be downloaded from https://doi.org/10.6084/m9.figshare.28302986.

## Conclusions

Here we report the completed genomes of 15 *S. sonnei* isolates, representing epidemiologically relevant and phylogenetically distinct genotypes. Both bacterial isolates and complete reference genomes are made publicly available to support future experimental and computational research on *S. sonnei*, which is rapidly emerging as the dominant agent of shigellosis. The in-depth characterisation of complete genome sequences presented here highlights evidence for ongoing adaptive evolution in *S. sonnei*, a topic that deserves further functional testing.

## Supplementary legends

**Figure S1. Clinker gene cluster comparison of Tn7/Int2 and *Shigella* resistance loci variants. A)** Gene cluster comparison of the Tn7 / Int2 region in representative clade 3.6 and 3.7 genomes. **B)** Gene cluster comparison of the *Shigella* resistance loci (SRL) pathogenicity island in lineages 2.1 and 3.4.1. Homologous regions are linked through the black/grey bars which also indicate the percentage identity. AMR = antimicrobial resistance.

**Figure S2. Whole genome alignment of *S. sonnei* chromosomal sequences using progressiveMauve.** Coloured blocks represent locally colinear blocks of sequence homology with respect to the reference genome (Lineage 1.5 in this case, indicated by (R)). Inverted regions are shown on the bottom strand. Position 1 corresponds to *fabB* in all genomes. Blank regions indicate regions with a reduced average sequence homology. Numbers at the top indicate chromosomal position. Mauve alignments can be downloaded from https://doi.org/10.6084/m9.figshare.28302986.

**Figure S3. Whole genome alignment of *S. sonnei* pINV sequences using progressiveMauve.** Coloured blocks represent locally colinear blocks of sequence homology, with respect to the reference genome (Lineage 1 in this case, indicated by (R)). Position 1 corresponds to *repA* in all genomes. Inverted regions are shown on the bottom strand. Blank regions indicate regions with an overall reduced average sequence homology. Mauve alignments can be downloaded from https://doi.org/10.6084/m9.figshare.28302986.

**Table S1. Summary of bacterial isolates used here.** This represents a more comprehensive version of Table 1. ^1^Genotypes were predicted using the *S. sonnei* genotyping framework implemented in Mykrobe, ^2^CDSs and tRNAs were predicted using Bakta. *Features of the completed genome of the classical laboratory strain, 53G, are included for comparative purposes. ONT, Oxford Nanopore Technologies. *tbd*, to be determined.

**Table S2. Summary of plasmid content in *S. sonnei* assemblies presented here.** Plasmids were replicon and MOB typed using the MobTyper function in MobSUITE. The top line of each row indicates plasmid size, and the bottom row indicates the accession of the nearest Mash neighbour. For MOBP and MOBQ, no replicon types were detected and for ‘No-MOB’ no replicon or MOB-types were identified. Plasmids of the completed genome of Lineage 2.8 are included. *denotes plasmids which encode antimicrobial resistance determinants.

**Table S3. Summary of plasmid-encoded colicins.** Colicins were identified using ABRicate and the full colicin database created previously [39]. This table directly corresponds to Table 2. If annotated, genes are indicated on the top row, with the protein product in brackets. The first column (IncFIA/IncFIC) represents pINV.

**Table S4. Summary of plasmid-encoded antimicrobial resistance determinants.** AMR determinants were identified using AMRfinder. This table directly corresponds to Table 2. The first column (IncFIA/IncFIC) represents pINV.

**Table S5. Summary of metabolic reactions predicted in each *S. sonnei* genome via CarveMe.** 1 indicates presence of the reaction, 0 indicates absence. Details of each reaction can be found in the xml model files at https://doi.org/10.6084/m9.figshare.28302986.

## Acknowledgements

We thank Mostowy and Holt lab members for helpful discussions.

Funding: SLM was supported by a Biotechnology and Biological Sciences Research Council LIDo Ph.D. studentship (BB/T008709/1). Research in the SM laboratory is supported by a Wellcome Trust Senior Research Fellowship (206444/Z/17/Z), European Research Council Consolidator Grant (772853 - ENTRAPMENT), and Wellcome Discovery Award (226644/Z/22/Z).

## References

1. Khalil, I.A., et al., Morbidity and mortality due to shigella and enterotoxigenic Escherichia coli diarrhoea: the Global Burden of Disease Study 1990-2016. The Lancet Infectious Diseases, 2018. 18(11): p. 1229–1240.

2. Sansonetti, P.J., D.J. Kopecko, and S.B. Formal, Involvement of a plasmid in the invasive ability of Shigella flexneri. Infection and Immunity, 1982. 35(3): p. 852–60.

3. Lan, R. and P.R. Reeves, Escherichia coli in disguise: molecular origins of Shigella. Microbes and infection, 2002. 4(11): p. 1125–1132.

4. Day, W.A., Jr., R.E. Fernández, and A.T. Maurelli, Pathoadaptive mutations that enhance virulence: genetic organization of the cadA regions of Shigella spp. Infection and Immunity, 2001. 69(12): p. 7471–80.

5. Yang, F., et al., Genome dynamics and diversity of Shigella species, the etiologic agents of bacillary dysentery. Nucleic Acids Research, 2005. 33(19): p. 6445–6458.

6. Parkhill, J., et al., Comparative analysis of the genome sequences of Bordetella pertussis, Bordetella parapertussis and Bordetella bronchiseptica. Nature Genetics, 2003. 35(1): p. 32–40.

7. Oré, N., et al., The decaying genome of Mycobacterium leprae. Leprosy Review, 2001. 72(4): p. 387–398.

8. Holt, K.E., et al., High-throughput sequencing provides insights into genome variation and evolution in Salmonella Typhi. Nature Genetics, 2008. 40(8): p. 987–993.

9. Zaghloul, L., et al., The distribution of insertion sequences in the genome of Shigella flexneri strain 2457T. FEMS Microbiology Letters, 2007. 277(2): p. 197–204.

10. Hawkey, J., et al., Impact of insertion sequences on convergent evolution of Shigella species. PLoS Genetics, 2020(1553-7404).

11. Holt, K.E., et al., Shigella sonnei genome sequencing and phylogenetic analysis indicate recent global dissemination from Europe. Nature Genetics, 2012. 44(9): p. 1056–9.

12. Thompson, C.N., P.T. Duy, and S. Baker, The Rising Dominance of Shigella sonnei: An Intercontinental Shift in the Etiology of Bacillary Dysentery. PLoS Neglected Tropical Diseases, 2015. 9(6): p. e0003708.

13. Holt, K.E., et al., Tracking the establishment of local endemic populations of an emergent enteric pathogen. Proceedings of the National Academy of Sciences, 2013. 110(43): p. 17522–17527.

14. Mason, L.C.E., et al., The evolution and international spread of extensively drug resistant Shigella sonnei. Nature Communications, 2023. 14(1): p. 1983.

15. Hawkey, J., et al., Global population structure and genotyping framework for genomic surveillance of the major dysentery pathogen, Shigella sonnei. Nature Communications, 2021. 12(1): p. 2684.

16. Baker, K.S., et al., Whole genome sequencing of Shigella sonnei through PulseNet Latin America and Caribbean: advancing global surveillance of foodborne illnesses. Clinical Microbiology and Infection, 2017. 23(11): p. 845–853.

17. Chung The, H., et al., Evolutionary histories and antimicrobial resistance in Shigella flexneri and Shigella sonnei in Southeast Asia. Communications Biology, 2021. 4(1): p. 353.

18. Martyn, J.E., et al., Maintenance of the Shigella sonnei Virulence Plasmid Is Dependent on Its Repertoire and Amino Acid Sequence of Toxin-Antitoxin Systems. Journal of Bacteriology, 2022. 204(3): p. e0051921.

19. Miles, S.L., K.E. Holt, and S. Mostowy, Recent advances in modelling Shigella infection. Trends in Microbiology, 2024. 32(9): p. 917–924.

20. Bardsley, M., et al., Persistent Transmission of Shigellosis in England Is Associated with a Recently Emerged Multidrug-Resistant Strain of Shigella sonnei. Journal of Clinical Microbiology, 2020. 58(4).

21. Lefèvre, S., et al., Rapid emergence of extensively drug-resistant Shigella sonnei in France. Nature Communications, 2023. 14(1): p. 462.

22. Duong, V.T., et al., No Clinical Benefit of Empirical Antimicrobial Therapy for Pediatric Diarrhea in a High-Usage, High-Resistance Setting. Clinical Infectious Diseases, 2017. 66(4): p. 504–511.

23. Qadri, F., et al., Congo red binding and salt aggregation as indicators of virulence in Shigella species. Journal of Clinical Microbiology, 1988. 26(7): p. 1343–1348.

24. Andrews, S. FastQC: A Quality Control Tool for High Throughput Sequence Data [Online]. 2010; Available from: http://www.bioinformatics.babraham.ac.uk/projects/fastqc/.

25. Wick, R.R. Filtlong. 2017 [cited 2024; Available from: https://github.com/rrwick/Filtlong.

26. Bolger, A.M., M. Lohse, and B. Usadel, Trimmomatic: a flexible trimmer for Illumina sequence data. Bioinformatics, 2014. 30(15): p. 2114–2120.

27. Bouras, G., et al., Hybracter: enabling scalable, automated, complete and accurate bacterial genome assemblies. Microbial Genomics, 2024. 10(5).

28. Kolmogorov, M., et al., Assembly of long, error-prone reads using repeat graphs. Nature Biotechnology, 2019. 37(5): p. 540–546.

29. Bouras, G., et al., How low can you go? Short-read polishing of Oxford Nanopore bacterial genome assemblies. Microbial Genomics, 2024. 10(6).

30. Zimin, A.V. and S.L. Salzberg, The genome polishing tool POLCA makes fast and accurate corrections in genome assemblies. PLoS Computational Biology, 2020. 16(6): p. e1007981.

31. Gurevich, A., et al., QUAST: quality assessment tool for genome assemblies. Bioinformatics, 2013. 29(8): p. 1072–5.

32. Parks, D.H., et al., CheckM: assessing the quality of microbial genomes recovered from isolates, single cells, and metagenomes. Genome Research, 2015. 25(7): p. 1043–55.

33. Shen, W., et al., SeqKit: A Cross-Platform and Ultrafast Toolkit for FASTA/Q File Manipulation. PLoS ONE, 2016. 11(10): p. e0163962.

34. Schwengers, O., et al., Bakta: rapid and standardized annotation of bacterial genomes via alignment-free sequence identification. Microbial Genomics, 2021. 7(11).

35. Hunt, M., et al., Antibiotic resistance prediction for Mycobacterium tuberculosis from genome sequence data with Mykrobe. Wellcome Open Research, 2019. 4: p. 191.

36. Robertson, J. and J.H.E. Nash, MOB-suite: software tools for clustering, reconstruction and typing of plasmids from draft assemblies. Microbial Genomics, 2018. 4(8).

37. Feldgarden, M., et al., AMRFinderPlus and the Reference Gene Catalog facilitate examination of the genomic links among antimicrobial resistance, stress response, and virulence. Scientific Reports, 2021. 11(1): p. 12728.

38. Seemann, T. Abricate. 2016 [cited 2022; Available from: https://github.com/tseemann/abricate.

39. De Silva, P.M., et al., Escherichia coli killing by epidemiologically successful sublineages of Shigella sonnei is mediated by colicins. EBioMedicine, 2023. 97: p. 104822.

40. Xie, Z. and H. Tang, ISEScan: automated identification of insertion sequence elements in prokaryotic genomes. Bioinformatics, 2017. 33(21): p. 3340–3347.

41. Tatusova, T., et al., NCBI Prokaryotic Genome Annotation Pipeline. Nucleic Acids Research, 2016. 44(14): p. 6614–24.

42. Tonkin-Hill, G., et al., Producing polished prokaryotic pangenomes with the Panaroo pipeline. Genome Biology, 2020. 21(1): p. 180.

43. Katoh, K. and D.M. Standley, MAFFT multiple sequence alignment software version 7: improvements in performance and usability. Molecular Biology and Evolution, 2013. 30(4): p. 772–80.

44. Price, M.N., P.S. Dehal, and A.P. Arkin, FastTree 2--approximately maximum-likelihood trees for large alignments. PLoS One, 2010. 5(3): p. e9490.

45. Hadfield, J., et al., Phandango: an interactive viewer for bacterial population genomics. Bioinformatics, 2017. 34(2): p. 292–293.

46. Thomas, P.D., et al., PANTHER: Making genome-scale phylogenetics accessible to all. Protein Science, 2022. 31(1): p. 8–22.

47. Darling, A.E., B. Mau, and N.T. Perna, progressiveMauve: multiple genome alignment with gene gain, loss and rearrangement. PLoS One, 2010. 5(6): p. e11147.

48. Gilchrist, C.L.M. and Y.-H. Chooi, *clinker &* clustermap.js: automatic generation of gene cluster comparison figures. Bioinformatics, 2021. 37(16): p. 2473–2475.

49. Machado, D., et al., Fast automated reconstruction of genome-scale metabolic models for microbial species and communities. Nucleic Acids Research, 2018. 46(15): p. 7542–7553.

50. Vezina, B., et al., Bactabolize is a tool for high-throughput generation of bacterial strain-specific metabolic models. eLife, 2023. 12: p. RP87406.

51. Calcuttawala, F., et al., Characterization of E-type colicinogenic plasmids from Shigella sonnei. FEMS Microbiology Letters, 2017. 364(7).

52. Chung The, H., et al., South Asia as a Reservoir for the Global Spread of Ciprofloxacin-Resistant Shigella sonnei: A Cross-Sectional Study. PLoS Medicine 2016. 13(8): p. e1002055.

53. Chung The, H., et al., Dissecting the molecular evolution of fluoroquinolone-resistant Shigella sonnei. Nature Communications, 2019. 10(1): p. 4828.

54. Baker, K.S., et al., Genomic epidemiology of Shigella in the United Kingdom shows transmission of pathogen sublineages and determinants of antimicrobial resistance. Scientific Reports, 2018. 8(1): p. 7389.

55. Turner, S.A., et al., Molecular epidemiology of the SRL pathogenicity island. Antimicrobial Agents and Chemotherapy, 2003. 47(2): p. 727–34.

56. Siguier, P., E. Gourbeyre, and M. Chandler, Bacterial insertion sequences: their genomic impact and diversity. FEMS Microbiol Reviews, 2014. 38(5): p. 865–91.

57. Seferbekova, Z., et al., High Rates of Genome Rearrangements and Pathogenicity of Shigella spp. Frontiers in Microbiology, 2021. 12: p. 628622.

58. Song, S. and C. Park, Organization and regulation of the D-xylose operons in Escherichia coli K-12: XylR acts as a transcriptional activator. Journal of Bacteriology, 1997. 179(22): p. 7025–32.

59. Bravo, V., et al., Distinct mutations led to inactivation of type 1 fimbriae expression in Shigella spp. PLoS One, 2015. 10(3): p. e0121785.

60. West, P.T., R.B. Chanin, and A.S. Bhatt, From genome structure to function: insights into structural variation in microbiology. Current Opinion in Microbiology, 2022. 69: p. 102192.

61. Payne, D., M. O’Reilly, and D. Williamson, The K88 fimbrial adhesin of enterotoxigenic Escherichia coli binds to beta 1-linked galactosyl residues in glycosphingolipids. Infection and Immunity, 1993. 61(9): p. 3673–7.

62. Sansonetti, P.J., D.J. Kopecko, and S.B. Formal, Shigella sonnei plasmids: evidence that a large plasmid is necessary for virulence. Infection and Immunity, 1981. 34(1): p. 75–83.

63. Prosseda, G., et al., Shedding of genes that interfere with the pathogenic lifestyle: the Shigella model. Research in Microbiology, 2012. 163(6): p. 399–406.

64. Rezwan, F., R. Lan, and P.R. Reeves, Molecular basis of the indole-negative reaction in Shigella strains: extensive damages to the tna operon by insertion sequences. Journal of Bacteriology, 2004. 186(21): p. 7460–5.

65. Thanh Duy, P., et al., Commensal Escherichia coli are a reservoir for the transfer of XDR plasmids into epidemic fluoroquinolone-resistant Shigella sonnei. Nature Microbiology, 2020. 5(2): p. 256–264.

