## Supplementary figures and images for "A public resource of 15 genomically characterised representative strains of *Shigella sonnei*"

### Supplementary figures

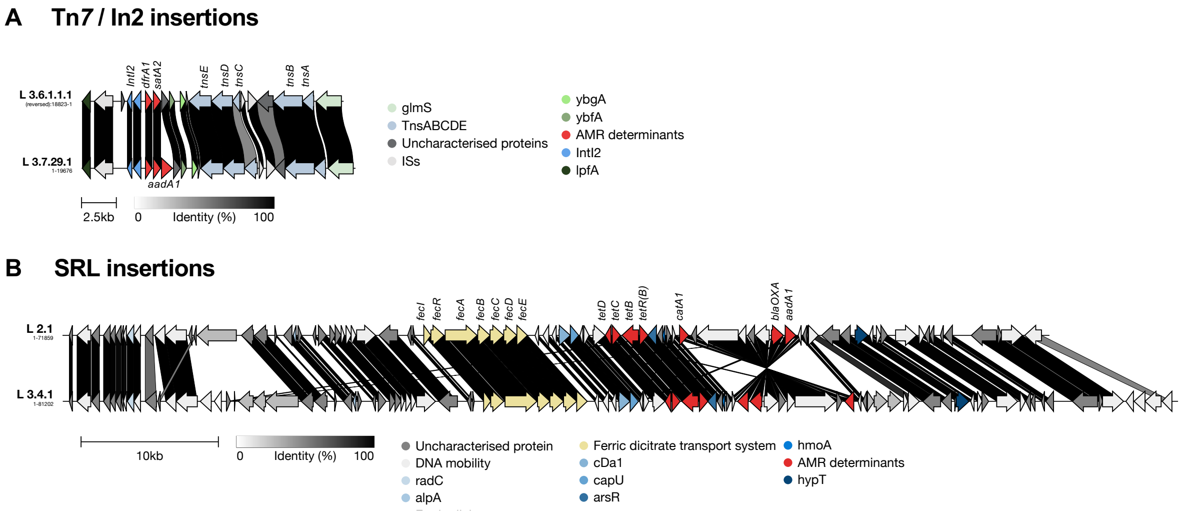

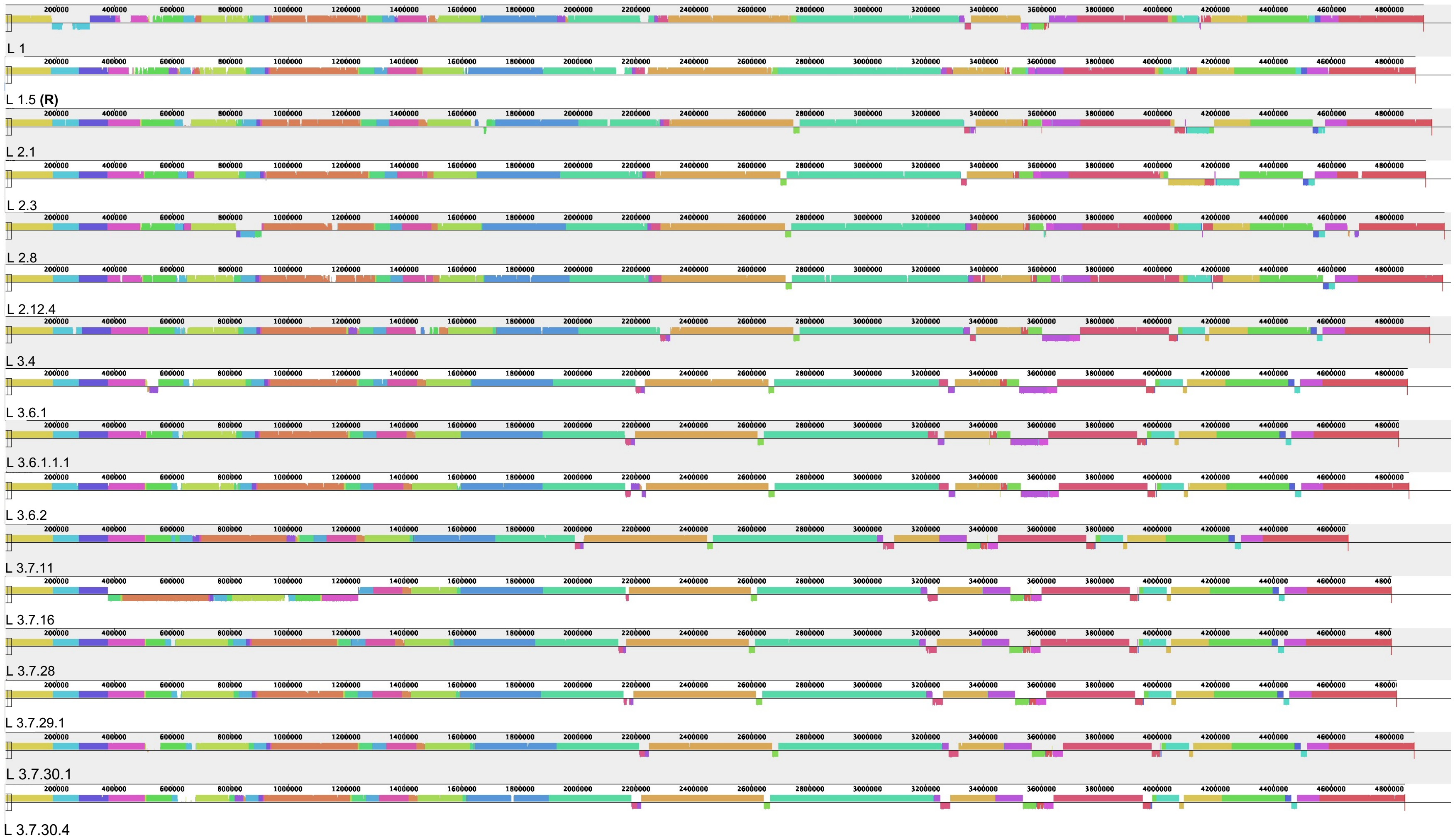

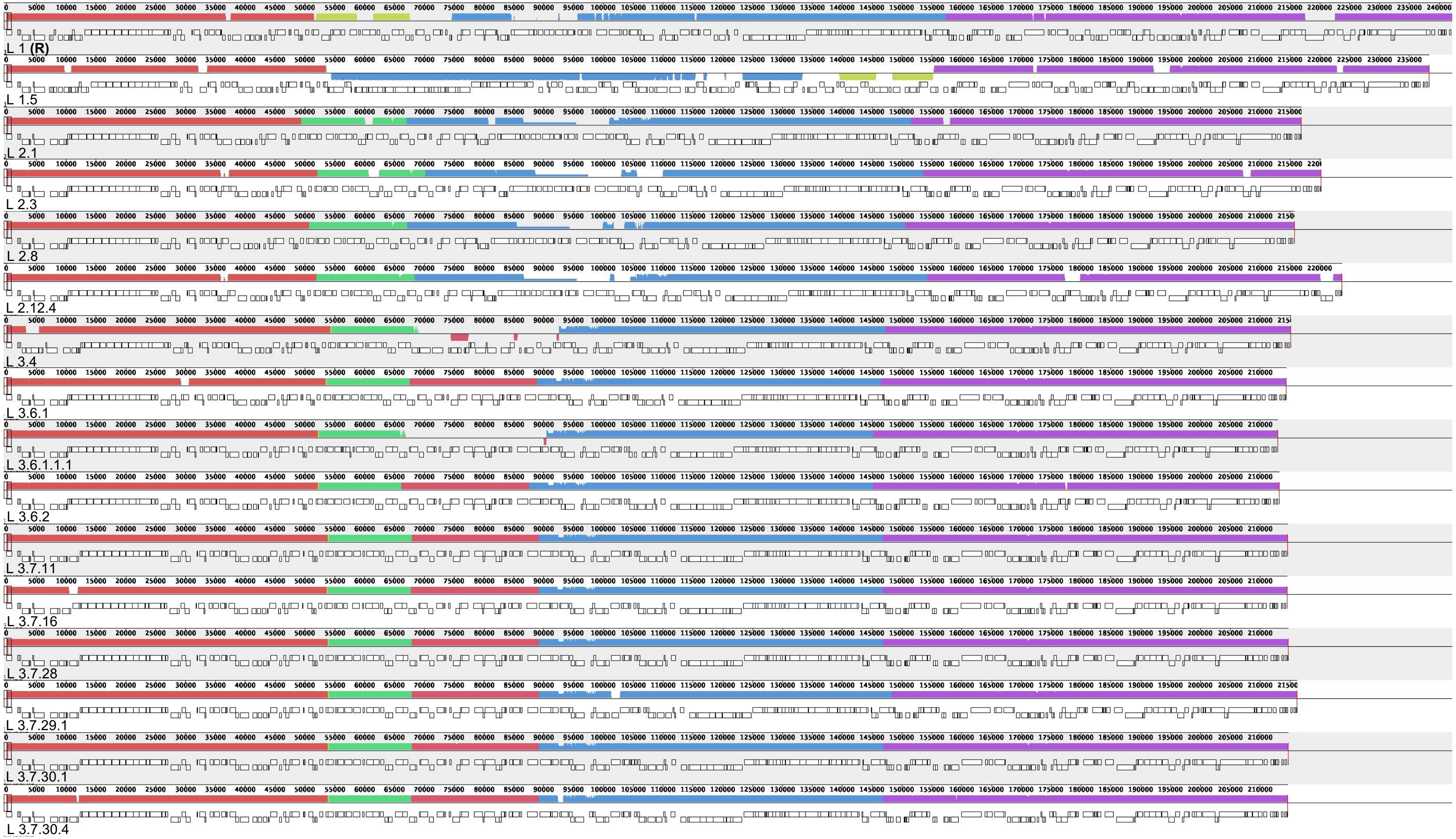
