## Supplementary files for "A public resource of 15 genomically characterised representative strains of *Shigella sonnei*"

**Table S1**

| Strain ID | Genotype[1] | Genome size (bp) | N50 | GC (%) | ONT depth | CDSs[2] | tRNAs | rRNAs | Contigs | Chromosome size (bp) | pINV size (bp) | Other plasmids | Complete-ness[3] | Contamin a-tion[3] | Genbank accession | Culture collection accession | Year of isolation | Country of isolation | Citati on |
|---|---|---|---|---|---|---|---|---|---|---|---|---|---|---|---|---|---|---|---|
| **201809330** | 1 | 5389660 | 4916714 | 50.8 | 61x | 5428 | 97 | 22 | 7 | 4916714 | 242731 | 5 | 99.54 | 0.26 | *tbd* | CIP112510 | 2018 | France | [21] |
| **391324** | 1.5 | 5260824 | 4888493 | 50.9 | 59x | 5255 | 97 | 22 | 4 | 4888493 | 238313 | 2 | 99.54 | 0.16 | *tbd* | *tbd* | 2017 | UK | N/A |
| **356538** | 2.1 | 5170588 | 4946399 | 50.8 | 76x | 5176 | 97 | 22 | 4 | 4946399 | 216858 | 2 | 99.56 | 0.11 | *tbd* | *tbd* | 2017 | UK | [14] |
| **830292** | 2.3 | 5150027 | 4924717 | 50.8 | 60x | 5166 | 96 | 22 | 3 | 4924717 | 220157 | 1 | 98.98 | 0.11 | *tbd* | *tbd* | 2019 | UK | [14] |
| **373220** | 2.12.4 | 5289501 | 4983563 | 50.8 | 47x | 5306 | 93 | 22 | 4 | 4983563 | 223662 | 2 | 99.6 | 0.11 | *tbd* | *tbd* | 2017 | UK | [14] |
| **590907** | 3.4.1 | 5174681 | 4940164 | 50.8 | 74x | 5229 | 97 | 22 | 7 | 4940164 | 215149 | 5 | 99.6 | 0.11 | *tbd* | *tbd* | 2018 | UK | [14] |
| **623218** | 3.6.1 | 5112917 | 4862817 | 50.8 | 68x | 5154 | 97 | 22 | 9 | 4862817 | 214316 | 7 | 99.6 | 0.26 | *tbd* | *tbd* | 2018 | UK | [14] |
| **02_1157** | 3.6.1.1.1 | 5073758 | 4832017 | 50.8 | 86x | 5110 | 96 | 22 | 5 | 4832017 | 212976 | 3 | 99.6 | 0.11 | *tbd* | *tbd* | 2014 | Vietnam | [52] |
| **642321** | 3.6.2 | 5100173 | 4866958 | 50.8 | 43x | 5135 | 97 | 22 | 6 | 4866958 | 213229 | 4 | 99.6 | 0.11 | *tbd* | *tbd* | 2018 | UK | [14] |
| **633497** | 3.7.11 | 4986415 | 4657310 | 50.8 | 46x | 5046 | 93 | 22 | 7 | 4657310 | 214646 | 5 | 97.74 | 0.11 | *tbd* | *tbd* | 2018 | UK | [14] |
| **598955** | 3.7.16 | 5039937 | 4807305 | 50.8 | 54x | 5067 | 98 | 22 | 7 | 4807305 | 214536 | 5 | 99.6 | 0.11 | *tbd* | *tbd* | 2018 | UK | [14] |
| **618335** | 3.7.28 | 5138354 | 4806102 | 50.7 | 61x | 5177 | 98 | 22 | 6 | 4806102 | 214666 | 4 | 99.6 | 0.58 | *tbd* | *tbd* | 2018 | UK | [14] |
| **03_0142** | 3.7.29.1.4 | 5156461 | 4824935 | 50.8 | 86x | 5182 | 96 | 22 | 9 | 4824935 | 214579 | 7 | 99.6 | 0.11 | *tbd* | *tbd* | 2014 | Vietnam | [65] |
| **627346** | 3.7.30.1 | 5122274 | 4884953 | 50.8 | 42x | 5168 | 100 | 22 | 6 | 4884953 | 216152 | 4 | 99.6 | 0.11 | *tbd* | *tbd* | 2018 | UK | [14] |
| **381259** | 3.7.30.4.1 | 5260132 | 4933886 | 50.7 | 58x | 5322 | 91 | 22 | 8 | 4933886 | 214700 | 6 | 99.6 | 0.11 | *tbd* | *tbd* | 2017 | UK | [14] |
| **53G*** | 2.8 | 5220473 | 4988504 | 50.7 | NA | 5248 | 96 | 22 | 5 | 4988504 | 215774 | 3 | NA | NA | *tbd* | *tbd* | 1954 | Japan | [11] |

**Table S2.**

| Genotype | IncFIA/<br>IncFIC | MOB$_F$/<br>Col-E1-<br>like | Col(BS512) | Col156/<br>MOB$_Q$ | Col(MG828) | IncI2/<br>MOB$_P$ | IncFIA,IncFII/<br>MOB$_F$,MOB$_P$ | IncK2/Z/<br>MOB$_P$ | IncI1/B/O<br>/MOB$_P$ | IncFIB | IncX1/MOB$_P$ | IncI-<br>gamma<br>/K1/MOB$_P$ | MOB$^P$ | MOB$_Q$ | No-MOB |
|---|---|---|---|---|---|---|---|---|---|---|---|---|---|---|---|
| **1** | 242731 bp*<br>***HE616529*** | - | - | - | - | 65348 bp*<br>***CP028155***<br>43917 bp*<br>***LT985310*** | - | - | 102906 bp*<br>***CP041565*** | - | - | - | 6888 bp<br>***CP014198*** | 11156 bp<br>***CP023649*** | - |
| **1.5** | 238313 bp<br>***CP000039*** | - | - | - | - | - | 125562 bp<br>***CP001065*** | - | - | - | - | - | - | 7084 bp<br>***CP023649*** | - |
| **2.1** | 216858 bp<br>***HE616529*** | - | - | - | - | - | - | - | - | - | - | - | - | 4074 bp<br>***CP018208*** | 3257 bp<br>***CP019023*** |
| **2.3** | 220157 bp<br>***HE616529*** | - | - | - | - | - | - | - | - | - | - | - | 5153 bp<br>***HE616530*** | - | - |
| **2.8** | 215774 bp<br>***HE616529*** | 5153 bp<br>***CP019897*** | 2089 bp<br>***HE616531*** | - | - | - | - | - | - | - | - | - | - | - | 8953 bp<br>***HE616532*** |
| **2.12.4** | 223662 bp<br>***CP023646*** | - | - | - | - | - | 77123 bp*<br>***AP014877*** | - | - | - | - | - | 5153 bp<br>***HE616530*** | - | - |
| **3.4.1** | 215149 bp<br>***CP023646*** | - | 2088 bp<br>***CP033398*** | 5114 bp<br>***CP019693*** | - | - | - | - | - | - | - | - | 2717 bp<br>***CP038000*** | 6750 bp<br>***DQ916145*** | 2699 bp*<br>***CP039609*** |
| **3.6.1** | 214316 bp<br>***CP023646*** | 2690 bp<br>***CP038000*** | - | 6015 bp<br>***KP970685***<br>5114 bp<br>***CP019693*** | 1549 bp<br>***CP003038*** | - | - | - | - | - | - | - | 7939 bp<br>***KU932034*** | 4076 bp<br>***CP011140*** | 8401 bp*<br>***CP034068*** |
| **3.6.1.1.1** | 212976 bp<br>***CP023646*** | - | 2089 bp<br>***CP030115*** | - | - | - | - | - | - | - | - | - | 2690 bp<br>***CP038000*** | 4878 bp<br>***KP970685***<br>4269 bp<br>***CP019693*** | 8401 bp*<br>***CP034068***<br>3787 bp<br>***CP019138***<br>2651 bp<br>***KX618698*** |
| **3.6.2** | 213229 bp<br>***CP023646*** | 2690 bp<br>***CP038000*** | 2089 bp<br>***CP030115*** | 5114 bp<br>***CP019693*** | - | - | - | - | - | - | - | - | - | - | 10093 bp*<br>***CP034068*** |
| **3.7.11** | 214646 bp<br>***CP016533*** | 2690 bp<br>***CP038000*** | 2101 bp<br>***CP023648*** | 5114 bp<br>***CP019693*** | - | - | - | - | - | - | - | 97999 bp<br>***CP016533*** | 6555 bp<br>***NC_008488*** | - | - |
| **3.7.16** | 214536 bp<br>***CP023646*** | 2690 bp<br>***CP038000*** | 2089 bp<br>***CP023648*** | 5114 bp<br>***CP019693*** | 1459 bp<br>***CP023264*** | - | - | - | - | - | - | - | - | 6744 bp<br>***KY348421*** | - |
| **3.7.28** | 214666 bp<br>***CP023646*** | 5153 bp<br>***CP019897***<br>2690 bp<br>***CP019897*** | - | - | - | - | - | - | - | 108503<br>bp<br>***CP03406<br>6*** | - | - | 1240 bp<br>***CP018984*** | - | - |
| **3.7.29.1.4** | 214579 bp<br>***CP023646*** | 2690 bp<br>***CP023653***<br>- | 2101 bp<br>***CP023648*** | - | 1549 bp<br>***CP003038*** | - | - | 96229 bp<br>***CP026855*** | - | - | - | - | - | 6774 bp<br>***NC_022585*** | 4869 bp<br>***CP030008***<br>2735 bp<br>***NC_011405*** |
| **3.7.30.1** | 216152 bp<br>***CP023646*** | 5153 bp<br>***CP023650***<br>2690 bp<br>***CP038000*** | - | - | - | - | - | - | - | - | - | - | - | - | 8212 bp<br>***CP025235*** |
| **3.7.30.4.1** | 214700 bp<br>***CP023646*** | 7717 bp<br>***CP023650*** | 2101 bp<br>***CP023648*** | 5114 bp<br>***CP019693*** | - | - | - | 57356 bp<br>***CP018984*** | - | - | 35184 bp<br>***CP002111*** | - | - | 4074 bp<br>***LT985255*** | - |

**Table S3**

| Genotype | IncFIA/ IncFIC | $MOB_F$/Col-E1-like | Col(BS512) | Col156/ $MOB_Q$ | Col(MG828) | IncI2/ $MOB_P$ | IncFIA,IncFII/ $MOB_F$,$MOB_P$ | IncK2/Z /$MOB_P$ | IncI1/B/O /$MOB_P$ | IncFIB | IncX1/ $MOB_P$ | IncI-gamma/K1 / $MOB_P$ | $MOB_P$ | $MOB_Q$ | No-MOB |
|---|---|---|---|---|---|---|---|---|---|---|---|---|---|---|---|
| **1** | - | - | - | - | - | - - | - | - | - | - | - | - | *cea* (Colicin-E1) | *ceaC* (Colicin-E3) | - |
| **1.5** | - | - | - | - | - | - | - | - | - | - | - | - | - | - | - |
| **2.1** | - | - | - | - | - | - | - | - | - | - | - | - | - | *ceaC* (Colicin-E3) | - |
| **2.3** | - | - | - | - | - | - | - | - | - | - | - | - | - | *ceaC* (Colicin-E3) | - |
| **2.8** | - | *cea* (Colicin-E1) | - | - | - | - | - | - | - | - | - | - | - | - | - |
| **2.12.4** | - | - | - | - | - | - |  | - | - | - | - | - | - | *ceaC* (Colicin-E3) | - |
| **3.4.1** | - | - | - | - | - | - | - | - | - | - | - | - | - | *col* (Colicin-E9) | - |
| **3.6.1** | - | - | - | (Colicin-E3) - | - | - | - | - | - | - | - | - | - | - | - |
| **3.6.1.1.1** | - | - | - | - | - | - | - | - | - | - | - | - | - | (Colicin-E3) - | - - - |
| **3.6.2** | - | - | - | - | - | - | - | - | - | - | - | - | - | - | - |
| **3.7.11** | - | - | - | - | - | - | - | - | - | - | - | *cia* (Colicin Ia) | - | - | - |
| **3.7.16** | - | - | - | - | - | - | - | - | - | - | - | - | - | *col* (Colicin-E9) | - |
| **3.7.28** | - | *cea* (Colicin-E1) - | - | - | - | - | - | - | - | - | - | - | - | - | - |
| **3.7.29.1.4** | - | - - | - | - | - | - | - | - | - | - | - | - | - | (Colicin-E5) | - - |
| **3.7.30.1** | - | *cea* (Colicin-E1) - | - | - | - | - | - | - | - | - | - | - | - | - | - |
| **3.7.30.4.1** | - | (Colicin 10) | - | - | - | - | - | - | - | - | - | - | - | - | - |

**Table S4**

| Genotype | IncFIA/IncFIC | MOB$_F$/Col-E1-like | Col(BS512) | Col156/MOB$_Q$ | Col(MG828) | IncI2/MO BP | IncFIA,IncFII/ MOB$_F$,MO BP | IncK2/Z /MO BP | IncI1/B/O /MO BP | IncFIB | IncX1/MO BP | IncI-gamma/K1 /MO BP | MO BP | MOB$_Q$ | No-MOB |
|---|---|---|---|---|---|---|---|---|---|---|---|---|---|---|---|
| **1** | *blaTEM-1* | - | - | - | - | *blaTEM-1*<br>*blaTEM-1* | - | - | *aph(6)-Id*<br>*blaTEM-1*<br>*dfrA14*<br>*aph(3'')-Ib*<br>*sul2* | - | - | - | - | - | - |
| **1.5** | - | - | - | - | - | - | - | - | - | - | - | - | - | - | - |
| **2.1** | - | - | - | - | - | - | - | - | - | - | - | - | - | - | - |
| **2.3** | - | - | - | - | - | - | - | - | - | - | - | - | - | - | - |
| **2.8** | - | - | - | - | - | - | - | - | - | - | - | - | - | - | - |
| **2.12.4** | - | - | - | - | - | - | *aph(6)-Id*<br>*blaTEM-1*<br>*aph(3'')-Ib*<br>*sul2*<br>*tet(B)* | - | - | - | - | - | - | - | - |
| **3.4.1** | - | - | - | - | - | - | - | - | - | - | - | - | - | - | *qnrB19* |
| **3.6.1** | - | - | - | -<br>- | - | - | - | - | - | - | - | - | - | - | *aph(6)-Id*<br>*aph(3'')-Ib*<br>*sul2*<br>*tet(A)* |
| **3.6.1.1.1** | - | - | - | - | - | - | - | - | - | - | - | - | - | -<br>- | *aph(6)-Id*<br>*aph(3'')-Ib*<br>*sul2*<br>*tet(A)*<br>-<br>- |
| **3.6.2** | - | - | - | - | - | - | - | - | - | - | - | - | - | - | *aph(6)-Id*<br>*aph(3'')-Ib*<br>*sul2*<br>*tet(A)* |
| **3.7.11** | - | - | - | - | - | - | - | - | - | - | - | - | - | - | - |
| **3.7.16** | - | - | - | - | - | - | - | - | - | - | - | - | - | - | - |
| **3.7.28** | - | -<br>- | - | - | - | - | - | - | - | *dfrA51* | - | - | - | - | - |
| **3.7.29.1.2** | - | -<br>- | - | - | - | - | - | *aph(6)-Id*<br>*aph(3'')-Ib*<br>*sul2* | - | - | - | - | - | - | -<br>- |
| **3.7.30.1** | - | - | - | - | - | - | - | - | - | - | - | - | - | - | - |
| **3.7.30.4.1** | - | - | - | - | - | - | - | - | - | - | - | - | - | - | - |

**Supplementary material**

Gene list of structural variants too large to be inserted into Table 5.

**~ 130 kbp inversion in clades 3.4 and 3.6**

Hypothetical protein, *yhiM, yhiN, pitA, uspB, uspA, dtpB, rsmJ, prlC, rlmJ, gorA,* hypothetical protein, *arsR, arsB, arsC, arsR,* hypothetical protein, *tnp, istB,* hypothetical protein, *tnp, tnp, tnp, tnp, yhiS*, hypothetical protein, *slp , dctR, yhiD, hdeB, hdeA, hdeD, gadE,* hypothetical protein, *mdtE, mdtF,* hypothetical protein, *gadW, gadX, gadB, ccp , treF, tnp, yieL,* hypothetical protein, hypothetical protein, *cbrC,* hypothetical protein, hypothetical protein, *cbrB, yieH, adeP, chrR,* hypothetical protein, *yieE, yidZ, mdtL*

**~107 kbp translocation inversion in clade 3.7**

*mnmE, yidC, rnpA, rpmH, ysdD, dnaA, dnaN, recF, gyrB, yidB, yidA, yidX, dgoR, dgoK, dgoA, dgoD, dgoD, dgoT, cbrA, yidR, yidQ, ibpA, ibpB, yidE, yidP, glvC,* hypothetical protein, *celF, insQ, yidI, yidH, yidG, yidF, emrD, ysdE, tisB,* hypothetical protein, *ilvN, uhpA, uhpB, uhpC, uhpT, adeD, tnp*, hypothetical protein, *yicN, nepI, yicS, nlpA,* hypothetical protein, hypothetical protein, hypothetical protein, hypothetical protein, tnp, hypothetical protein, tnp, *tnp,* hypothetical protein, *tnp, iutA, iucD, iucC, iucB, iucA, shiF,* hypothetical protein, hypothetical protein, *tnp, tnp, tnp,* hypothetical protein, *tra5,* hypothetical protein, *tnp, tnp,* hypothetical protein, *tnp, istB,* hypothetical protein, *tnp, tnp, tnp, tnp, yhiS,* hypothetical protein, *slp , dctR, yhiD, hdeB, hdeA, hdeD, gadE,* hypothetical protein, *mdtE, mdtF,* hypothetical protein, *gadW, gadX, gadB, ccp , treF,* hypothetical protein, *tnp, yieL,* hypothetical protein, *tnp, cbrC,* hypothetical protein, hypothetical protein, *cbrB, yieH, adeP, chrR,* hypothetical protein, *yieE, yidZ, mdtL*
